# Lateral line hair cells integrate mechanical and chemical cues to orient navigation

**DOI:** 10.1101/2022.08.31.505989

**Authors:** Laura Desban, Julian Roussel, Olivier Mirat, François-Xavier Lejeune, Ludovic Keiser, Nicolas Michalski, Claire Wyart

**Author notes:** Co-corresponding authors: Claire Wyart, Laura Desban. These authors contributed equally to this work (Laura Desban, Julian Roussel).

## Abstract

The lateral line is a superficial sensory system responding to environmental hydrodynamic changes to orient locomotion of aquatic vertebrate species. Whether this system also detects chemical cues is unknown. We find that zebrafish lateral line hair cells express numerous chemoreceptors, including ionotropic receptors for serotonin. We show that the serotonin enriched in skin neuroepithelial cells is released upon injury and that environmental serotonin activates lateral line hair cells. We show that larval zebrafish exposed to serotonin in their environment rely on the lateral line to swim fast and away. These results uncover the sensory versatility of lateral line hair cells and how these properties modulate navigation in response to environmental stimuli.

## Introduction

All cartilaginous and bony fish exhibit a superficial sensory system, called the lateral line, which is exposed to the surrounding water and covers their entire body surface (*1*). The lateral line relies on hair cells, highly similar to the ones responsible for acousto-vestibular functions in the ear, organized in discrete organs called neuromasts (*1*). Since its discovery, substantial evidence has shown that the lateral line detects mechanical stimuli associated with hydrodynamic changes in the surrounding water, thereby conveying the sense of “touch-at-a-distance” (*2*). In this way, the lateral line is a major sensory system in fish that contributes to complex locomotor behaviors such as schooling (*3, 4*), predation (*5*), and orientation to the large-scale flow field, referred to as rheotaxis (*6, 7*).

Given its spatial organization at the body surface, the lateral line would be ideally located to support the detection of chemicals and chemical gradients in the environment, a fundamental property for aquatic animals to find optimal living conditions. Although former studies on cochlear hair cells have reported the expression of putative chemoreceptors such as transient receptor potential (TRP) channels (*8–11*) and purinergic receptors (*12, 13*), they only investigated their contribution to mechanoelectrical transduction, thereby leaving the question of chemoreception unaddressed. To tackle this question, we investigated whether chemoreception occurs in the lateral line of larval zebrafish to sense chemicals in the surrounding water.

## Results

### Lateral line hair cells express numerous chemoreceptors

To probe the sensory repertoire of lateral line hair cells, we performed an unbiased transcriptome analysis of these cells. Due to the lack of transgenic lines specifically labelling hair cells of the lateral line and not of the inner ear, we turned to the vital dye Yo-Pro-1 (*14*) that selectively labels active, superficial hair cells (Fig. 1A). We bath-applied 3 days post-fertilization (dpf) zebrafish larvae with this dye and isolated Yo-Pro-1-positive and -negative cell fractions using fluorescent-activated cell sorting (FACS) (Fig. 1B).

**Fig. 1.**
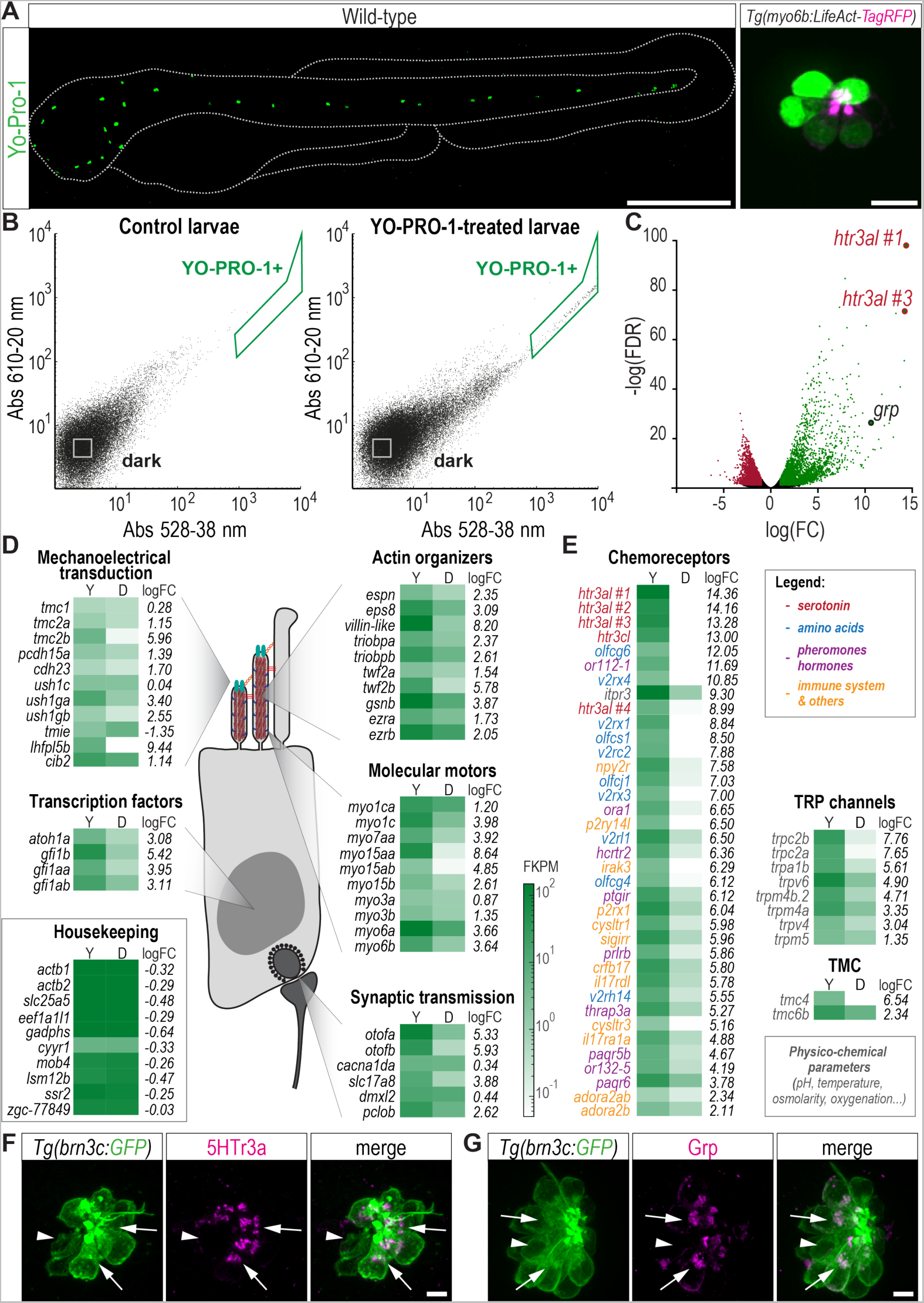
The transcriptome of lateral line hair cells unravels numerous chemosensory receptors. (**A**) Yo-Pro-1 specifically labels lateral line hair cells. Left, lateral view of a 3 dpf zebrafish larva after bath-application of Yo-Pro-1 (scale bar, 500 µm). Right, one neuromast showing Yo-Pro-1 labelling (green) in LifeAct-tagRFP-positive hair cells (magenta, actin) in a 3 dpf *Tg(myo6b:LifeAct-tagRFP)* larva (scale bar, 10 µm). (**B**) Illustrative FACS scatter plots from unlabeled (left) and Yo-Pro-1-labelled (right) sibling larvae. (**C**) Volcano plot showing transcripts down-(red dots) or up- (green dots) regulated in Yo-Pro-1-positive fractions. (**D**) Schematics of a hair cell. Transcripts for known hair cell genes are enriched in Yo-Pro-1-positive (Y) versus dark (D) fractions. FKPM, fragments per kilobase million. (**E**) List of 47 most enriched transcripts encoding for chemoreceptors, color-coded according to their ligand. (**F-G**) Validation of the expression and localization of the 5HTr3a receptor (**F,** arrows) and Grp peptide (**G**, arrows) in GFP-positive neuromast hair cells in a 5 dpf *Tg(brn3c:GFP)* larva. Arrowheads, non-expressing cells. Scale bars, 5 µm.

RNA sequencing of both fractions revealed 2522 up- and 1050 down-regulated transcripts in the Yo-Pro-1-positive population compared to the remainder of the tissue (Fig. 1C, and Data S1). The expression of numerous genes involved in landmark hair-cell functions such as mechanoelectrical transduction, hair bundle differentiation, structure and function of the ribbon synapse (Fig. 1D) confirmed that sorted Yo-Pro-1-positive fractions were successfully enriched in lateral line hair cells.

Our transcriptome analysis of lateral line hair cells also uncovered the expression of more than 45 novel chemoreceptors involved in the detection of a wide diversity of chemical cues (see shortlist of candidates with consistent strong enrichment across 5 biological replicates in Fig. 1E, and Data S1). Nineteen transcripts encode G protein-coupled receptors corresponding to either classical olfactory receptors, type C / vomeronasal receptors type 2 thought to detect amino-acids in zebrafish (*15*), or specific receptors for hormones and pheromones such as PGF2α (*16*). Twelve other transcripts encode receptors for various chemicals including neuropeptides, purine nucleotides and nucleosides, interleukins and leukotrienes. Another eight transcripts encode for members of the TRP family involved in detecting physicochemical parameters such as pH, temperature, and osmolarity (*17*), some of which had been previously identified in hair cells of the mammalian inner ear (*8–11*). Finally, and most strikingly, we found among the ten most enriched transcripts, five – *htr3al #1, htr3al #2, htr3al #3, hrt3al #4* and *htr3cl* – encoding isoforms of the non-selective ionotropic serotonin 3 receptor (5HT3r), suggesting an important modulation of lateral line hair cells by serotonin.

### A subset of hair cells expresses ionotropic serotonin receptors

To confirm the massive enrichment of transcripts for 5HTr3a receptors in lateral line hair cells as revealed by our transcriptomic analysis, we performed immunohistochemistry for 5HTr3a on whole-mounted 3 and 5 dpf *Tg(brn3c:GFP)* transgenic larvae where all hair cells are labelled with GFP (Fig. 1F). We used as a technical positive control the staining against a peptide found highly expressed in the lateral line, namely Grp, and whose transcripts were similarly enriched as *5htr3* (Fig. 1G). The 5HTr3a receptor was detected in 32.4 ± 16.9 % of hair cells at 3 days (76 cells imaged from 7 neuromasts in 4 larvae) and 69.9 ± 17.8 % at 5 days (75 cells imaged in 8 neuromasts in 5 larvae) where it was found in the subapical region, most likely corresponding to the Golgi transfer apparatus (*18*) (Fig. 1F). Other receptors such as the TRP channels Trpm4 and Trpa1, both of which had been previously identified in mouse inner ear hair cells (*8, 10*), were also detected by immunohistochemistry in a subset of lateral line hair cells (Fig. S1A-B). These results confirm that a fraction of lateral line hair cells express chemoreceptors, and could therefore respond to chemical cues. Additionally, along with confirming the presence of Trpm4 in stereocilia of inner ear hair cells (Fig. S1D,F), we uncovered strong expression of serotoninergic 5Htr3a receptors (Fig. S1C,E) in all cristae of the inner ear, suggesting that chemosensory properties may be common to different hair cell types in zebrafish.

### Environmental serotonin can be released in the water by injured conspecifics

The enrichment in ionotropic 5HTr3a receptors and their subcellular location in lateral line hair cells exposed at the surface of zebrafish skin suggest that these cells could detect serotonin from the surrounding water. We therefore asked whether and how serotonin could be released by conspecifics in the environment. We first performed immunohistochemistry for serotonin on whole-mounted larvae. Consistent with previous reports (*19–21*), we found that serotonin localizes to neurons of the pineal gland in the brain, taste receptor cells and enteroendocrine cells in the gut (Fig. 2A, see also Fig. 4C). We also observed the massive accumulation of serotonin in neuroepithelial cells (*22*) covering the entire body surface (Fig. 2A). Considering its accumulation in the skin, in direct contact with the environment, we tested whether serotonin could be released in the water following skin lesion (*23*). Measures with enzyme-linked immunosorbent assay (ELISA, see Methods) revealed no serotonin in the conditioned water of intact juvenile or adult fish (Fig. 2B, ‘ctrl’). In contrast, serotonin was present in solutions containing the whole-body extract of juveniles or adults (Fig. 2B, 8.3 ± 1.6 nM in juveniles, 3 replicates; 7.2 nM ± 2.5 nM in adults, 3 replicates) and skin extract (Fig. 2B, 3.1 ± 0.2 nM extracted from 1 adult fish ‘Skin 1’, 6 replicates; 16.6 nM ± 2.7 nM extracted from 2 adult fish ‘Skin 2’, 3 replicates), indicating that serotonin is released by injured fish in the environment and could therefore act as a potential cue of danger.

**Fig. 2.**
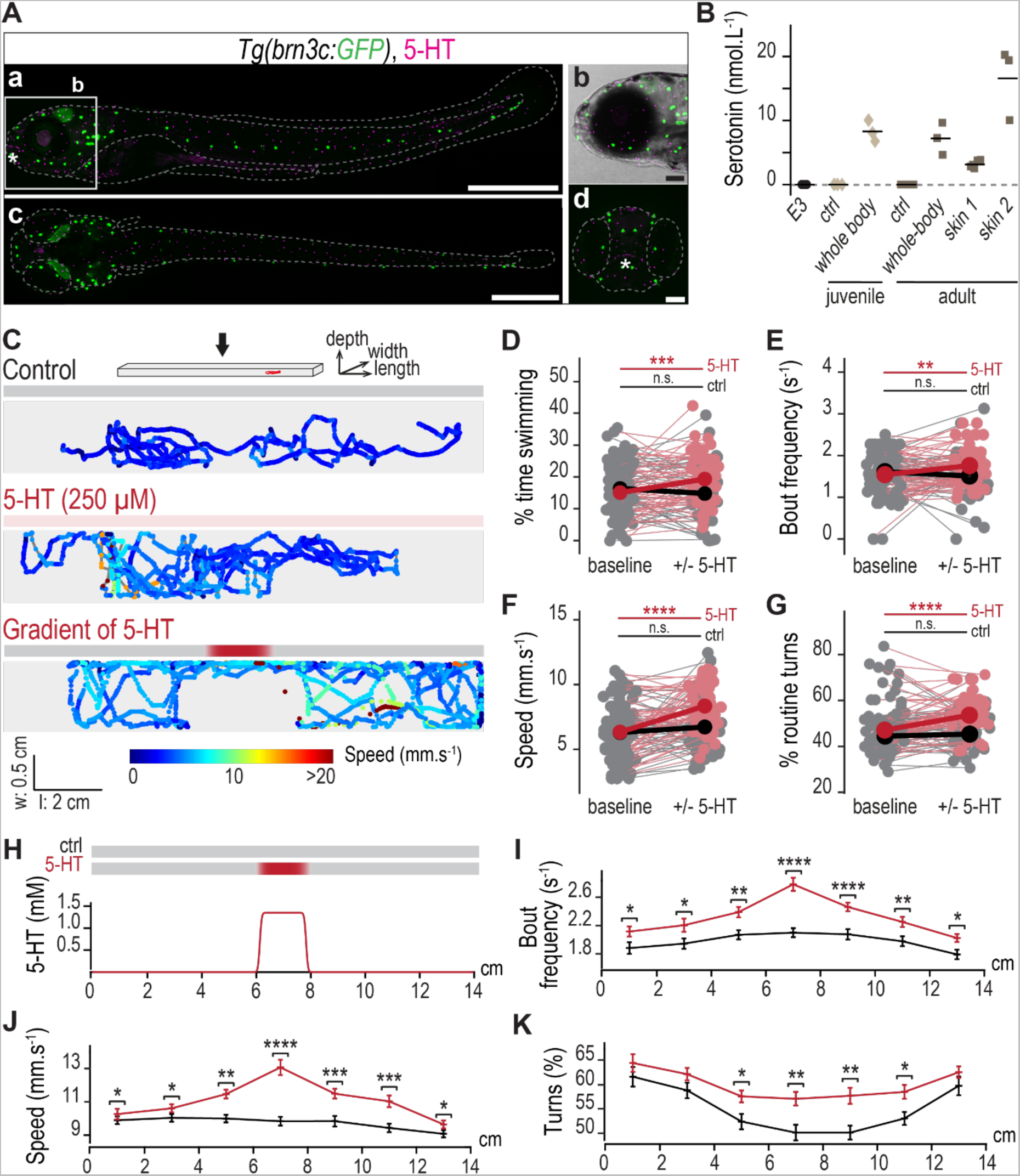
Environmental serotonin can be released by injured conspecifics and elicit avoidance behavior in larval zebrafish. (**A**) Neuroepithelial cells scattered on the skin that are in the vicinity of lateral line hair cells (GFP, green) are filled with serotonin (5-HT, magenta) in a typical 5 dpf larva: lateral (**a**, scale bar: 500 µm; and zoom in region **b**, scale bar: 100 µm), dorsal (**c**, scale bar: 500 µm), and frontal (**d**, scale bar: 50 µm) views. Asterisk, taste receptor cells expressing 5-HT. (**B**) Serotonin is found in adult zebrafish skin extract solution (skin extract from 1 fish “skin 1” – 6 replicates ; skin extract from 2 fish “skin 2”, 3 replicates) as well as in the whole-body extract solution of juveniles (3 replicates) and adults (3 replicates). Each point represents a biological replicate and the black bar the mean. (**C**) Typical trajectory of individual 5 dpf larvae recorded for 5 min either in control solution (top), homogenous serotonin (250 µM, middle) or in a steep gradient of serotonin (peak at 1mM, bottom). (**D-G**) Larvae exposed to homogeneous serotonin (magenta, N = 52 larvae) increase swimming and turns from baseline compared to siblings in control solution (grey, N = 47 larvae) with more time spent swimming (**D**), higher bout frequency (**E**), higher speed (**F**), and more turns (**G**) (type II Wald chi-square tests indicate a significant interaction effect between time and condition for percentage of time swimming ***p = 3x10^-4^, bout frequency **p = 0.001, speed ***p = 4x10^-4^ and percentage of routine turns **p = 0.003; statistics on the graph denote the results of post-hoc tests using Tukey’s method for the significant variation between baseline and test recordings). (**H-K**) A gradient of serotonin simulated in (**H**) locally affects navigation by increasing bout frequency (**I**), speed (**J**) and turning occurrence (**K**) (N = 87 larvae in control solution versus 89 in serotonin gradient; type II Wald chi-square tests indicate a significant interaction effect between position and condition for bout frequency **p = 1.3x10^-4^ and speed **p = 1.2x10^-3^; statistics on the graph denote the results of post-hoc tests using Tukey’s method comparing control and serotonin conditions at each position in the well). Note that Y scales do not systematically show the 0 in order to zoom on the group effect and variability across fish.

### Environmental serotonin elicits aversive behavior in larval zebrafish

To test the physiological relevance of serotonin release in water, we analyzed the behavior of freely-swimming larval zebrafish exposed to environmental serotonin (Fig. 2C, and Movie S1). Zebrafish larvae maintained in control solution retained consistent swimming over time (Fig. 2D-G). In contrast, larvae exposed to serotonin swam more (Fig. 2D-E, 22 % increase in time spent swimming and 14-% increase in bout frequency) and faster (Fig. 2F, 32 % increase in speed). Interestingly, the categorization of bouts into forward swims versus turns (see Methods) revealed that larvae also turned significantly more when exposed to serotonin (Fig. 2G, 11 % increase), suggesting an avoidance response to serotonin.

Next, we investigated whether zebrafish larvae could respond to a local source of serotonin. We introduced serotonin in the central region of swimming arenas and used simulations to show that the chemical slowly diffused over the time course of behavior recordings to form a sharp gradient with a peak concentration of 1 mM in the center of the arena (Fig. 2H, and Movie S2). As expected, zebrafish larvae navigating through the local gradient of serotonin swam more often and faster (Fig. 2I-J, and Movie S3). Supporting our hypothesis of an aversive response, larvae also turned significantly more when encountering serotonin (Fig. 2K). Together, these results indicate that zebrafish larvae respond to serotonin by swimming more often, faster and away from it.

### Lateral line hair cells respond to serotonin from the environment

Based on the fact that lateral line hair cells express 5HTr3a receptors, we asked whether these cells mediate the detection of environmental serotonin. We used a genetically-encoded calcium indicator GCaMP5G expressed under the control of the *brn3c* promoter (Fig. 3A) to monitor the response of lateral line hair cells upon pressure-application of serotonin in solution (Fig. 3B, and Movie S4). Our simulation shows that during the stimulus, an estimated maximum of ∼ 80 % of the initial 1 mM serotonin concentration in the pipet effectively reaches the cupula covering the neuromast hair cell bundle and rapidly decays in a few seconds (Fig. 3C, and Movie S5) enabling us to repeatedly test the transient exposure of lateral line hair cells to serotonin.

**Fig. 3.**
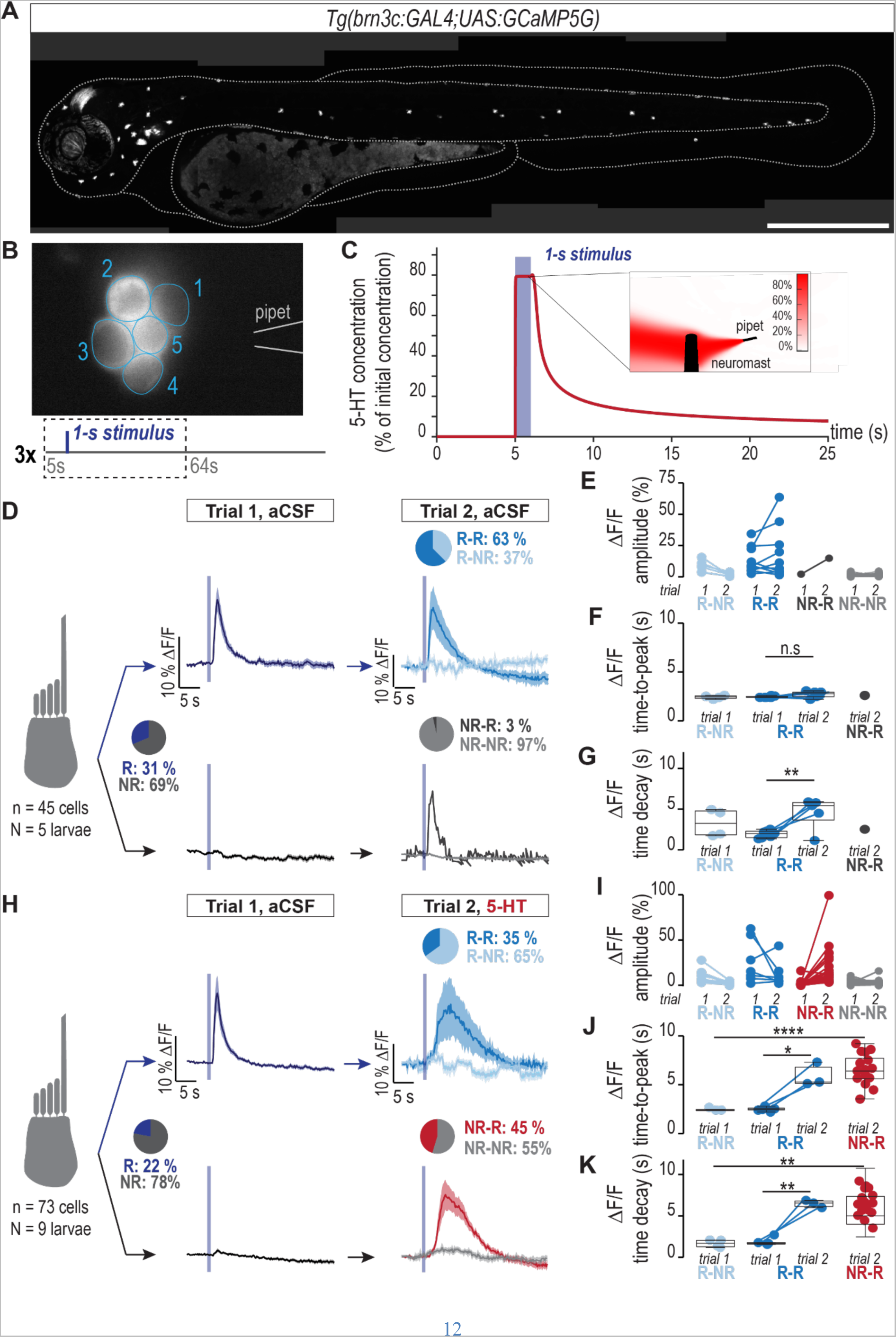
Environmental serotonin activates lateral line hair cells. (**A**) *Tg(brn3c:GAL4; UAS:GCaMP5G)* transgenic larva used to assess calcium activity of neuromast hair cells at 3 dpf (scale bar, 500 µm). (**B**) Schematics of the chemosensory assay: a glass pipet filled with the test solution is positioned at the vicinity of the hair bundle of one neuromast. GCaMP5G fluorescence intensity is monitored in each cell (regions of interest 1 to 5) while first tested with aCSF alone (Trial 1), then with aCSF containing serotonin using a new pipet (Trial 2). (**C**) Simulation of the variation over time of the concentration of serotonin, expressed as % of the initial concentration in the pipet (1 mM), that effectively reaches the neuromast cupula during and after the 1-s puff application. Inset, spatial distribution of the serotonin concentration at the end of the pressure-application (single time frame from Movie S5). (**D**) During a control experiment, hair cells were tested with aCSF in both trials (45 cells in 5 larvae). Among 31 % of cells responding during Trial 1 (“R”), 63 % remained responsive during Trial 2 (“R-R”). 97 % of non-responding cells during Trial 1 (“NR”) remained silent during Trial 2 (“NR-NR”). Traces, mean response across individual cells. Shade, standard error of the mean. Quantification of the ΔF/F amplitude (**E**), ΔF/F time-to-peak (**F**) and ΔF/F time decay (**G**) of mechanosensory responses depicted in **D** (paired t-tests, ^n.s.^p_time-to-peak_ = 0.13 and **p_decay_ 0.003). (**H**) 45 % of non-responding cells (73 tested cells in 9 larvae) during Trial 1 show a significant response to puffs of 1 mM serotonin (5-HT) during Trial 2 (“NR-R” in magenta). Quantification of the ΔF/F amplitude (**I**), ΔF/F time-to-peak (**J**) and ΔF/F time decay (**K**) of mechano- and chemo-sensory responses depicted in **H.** Compared to pure mechanosensory responses during Trial 1 (“R-NR”), mixed (“R-R”) or chemosensory only (“NR-R”) responses to serotonin show a significant increase in time-to-peak (paired t-tests to compare between trials for “R-R” cells: *p_time-to-peak_ = 0.033 and **p_decay_ = 0.003, and t-tests to compare between “R-NR” and “NR-R”: ****p_time-to-peak_ = 1x10^-5^ and **p_decay_ = 0.003).

We first investigated the reliability of the mechanical response upon repeated pressure-applications of a control solution of artificial cerebrospinal fluid (aCSF) using different pipettes with identical resistance and orientation (Fig. 3D, and Movie S6). Upon first aCSF application (“Trial 1 – aCSF”), we found that 31 % of all cells systematically responded (labeled “R”) to the mechanical stimulus (Fig. 3D, 14 cells out of 45), a result consistent with the orientation-based selectivity of lateral line hair cells (*24*). Out of the responding cells, two thirds (63 %) of them exhibited again a strong mechanical response when stimulated a second time with control aCSF (Fig. 3D, “Trial 2 – aCSF”, 9 cells out of 14 “R” cells, labeled “R-R”). In contrast, non-responding cells (labeled “NR”) during the first aCSF application remained robustly silent when subjected to a second pressure-application (Fig. 3D, 30 cells out of 31 “NR” cells, labeled “NR-NR”), demonstrating that the mechanosensory selectivity was consistent across trials. Note that the amplitude of the mechanical response varied within and across cells (Fig. 3E) with a significant habituation effect of time on the decay upon repeated stimulations (Fig. 3G, median of 2.0 s, 25^th^-75^th^ percentiles: 1.5-2.3 s in Trial 1, and 5.4 s with 25^th^-75^th^ percentiles: 3.7-5.8 s in Trial 2). Conversely, the time-to-peak of lateral line hair cell mechanical response remained consistent across cells and trials (Fig. 3F, median of 2.4 s in Trial 1 with 25^th^-75^th^ percentiles: 2.4-2.5 s, and 2.7 s in Trial 2 with 25^th^-75^th^ percentiles: 2.5-2.9 s).

Since non-responding cells remained unresponsive upon repeated mechanical stimulations with pipettes of same properties and orientation, we were able to identify non-responding cells (“NR”) in the first trial and test their response to serotonin pressure-applied during the second trial (Fig. 3H, and Movie S7). Consistent with our finding that only a fraction of lateral line hair cells express ionotropic serotonin 3 receptors at 3 dpf, 45 % of cells showed calcium transients in response to serotonin (Fig. 3H, “Trial 2 – 5-HT”, 26 cells out of 57, labeled “NR-R” in magenta), demonstrating that lateral line hair cells respond to serotonin in the surrounding water.

Although the response to serotonin was similar to the mechanical response in terms of amplitude (Fig. 3I, magenta), its kinematics were slower as reflected by a significant increase in median time-to-peak (Fig. 3J, median 6.4 s with 25^th^-75^th^ percentiles: 5.6-7.7 s, compared to 2.4 s with 25^th^-75^th^ percentiles: 2.4-2.5 s for the mechanosensory response) and in time decay (Fig. 3K, median 5.0 s with 25^th^-75^th^ percentiles: 4.0-7.3 s, compared to 2.0 s with 25^th^-75^th^ percentiles: 1.5-2.3 s for the mechanosensory response) (see also Movie S7). Interestingly, in cells responding to the mechanical stimulus in the first trial (“R” cells), pressure-application of serotonin also slowed down the kinematics of the mixt mechano/chemosensory response by significantly increasing on- and off-kinematics of mechanical activity (Fig. 3I-J, Trial 2 in “R-R” cells with median time to peak of 6.1 s with 25^th^-75^th^ percentiles: 5.0-8.3 s, and median time decay of 6.6 s with 25^th^-75^th^ percentiles: 5.0-8.3 s), suggesting that chemical activation by serotonin can modulate the mechanosensory-driven response of hair cells. Altogether, these results demonstrate that lateral line hair cells can integrate both chemical and mechanical cues, and that chemical signaling modulates the kinematics of hair cell mechanical response.

### Mechanosensory-dependent activity in lateral line hair cells sets basal levels of spontaneous swimming

We next investigated how lateral line hair cells contribute to the behavioral response to environmental serotonin (Fig. 4). Most studies investigating the role of the lateral line in behavior have focused on animals swimming in turbulent flows (*7, 25*). We first assessed the role of lateral line hair cells in the spontaneous locomotion of larval zebrafish swimming alone in still water. Zebrafish larvae swim in discrete bouts (*26, 27*) (Fig. 4A) interspaced by periods of quiescence, referred to as inter-bout intervals (IBIs), that follow a negative binomial distribution (*28*) (Fig. 4B). We performed cell-specific ablation of lateral line hair cells using two commonly used chemical treatments: waterborne copper sulfate (CuSO_4_) and BAPTA. CuSO_4_ is a known ototoxin that specifically eliminates neuromast hair cells (*29*) while BAPTA is a calcium chelator that disrupts tip links between stereocilia responsible for the gating of mechanoelectrical transduction channels upon hair bundle deflection, and thereby forbids mechanosensory-driven activity in a reversible manner without killing hair cells (*30*). As expected, exposure to CuSO_4_ induced the death of lateral line hair cells but left intact other superficial and / or sensory cells, including inner ear hair cells, neuroepithelial cells, and olfactory receptor neurons (Fig. 4C-D). Neuromast support cells, although partially disorganized due to the loss of neighboring hair cells within the same neuromast, also remained intact after treatment (Fig. 4D, middle panel). We did not observe any change in overall cell morphology and organization following BAPTA treatment (Fig. 4D, bottom panel) compared to untreated siblings (Fig. 4C-D, top panel). Quantification of hair cell numbers in inner ear structures and in neuromasts confirmed our observations (Fig. 4F) and validated the use of these two drugs to efficiently and specifically manipulate hair cell function.

We checked the effects of these drugs on sensorimotor integration involving the inner ear hair cells by testing acousto-vestibular escape responses. We found that 92 % of drug-treated larvae successfully performed acousto-vestibular escapes after CuSO_4_ (11 out of 12 fish, success rate of 60 %) or BAPTA (11 out of 12 fish, success rate of 53 %) treatment, similar to untreated siblings (12 out of 12 fish, success rate of 93 %) (Fig. 4G). We observed that treated larvae also responded to tactile stimuli during experimental handling, indicating that drug treatments did not affect inner ear hair cells and were not toxic for zebrafish larvae. We then assessed navigation in larvae with intact or impaired lateral line function by measuring fraction of time spent swimming, bout frequency and bout speed. Complete elimination of neuromast hair cells using CuSO_4_ drastically impacted spontaneous swimming (Fig. 4E, and Movie S8), observed as a 77 % decrease in time spent swimming (Fig. 4H), a 31 % decrease in bout frequency (Fig. 4I) and a 38 % decrease in speed (Fig. 4J). This result is consistent with previous findings in 1-week-old zebrafish larvae treated with either CuSO_4_ or neomycin, another ototoxic agent (*31*).

**Fig. 4.**
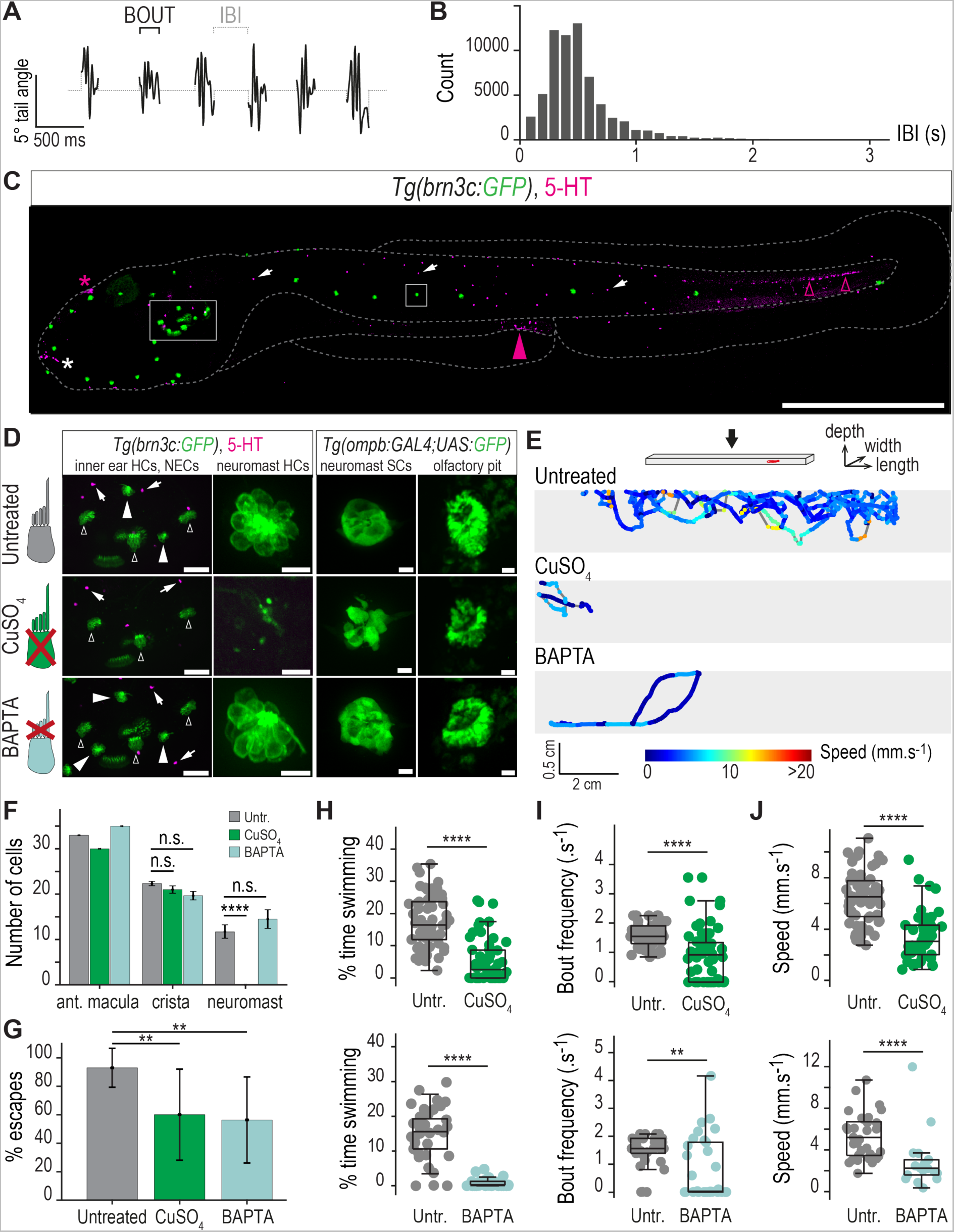
Mechanosensory-driven activity of lateral line hair cells sets basal levels of spontaneous swimming. (**A**) Tail angle over time illustrates the stereotyped swimming behavior of 5 dpf zebrafish larvae with discrete events of swim (bouts) interspaced by interbout intervals (IBIs). (**B**) IBIs follow a negative binomial distribution (n = 67,600 bouts). (**C**) Untreated 5 dpf *Tg(brn3c:GAL4; UAS:GFP)* larva with intact lateral line (square region) and inner ear (rectangular region) hair cells as revealed by immunostaining against GFP (green). Immunostaining against serotonin (5-HT, magenta) labels neuroepithelial cells on the skin (arrows), taste receptor cells (white asterisk), the pineal gland (magenta asterisk), enteroendocrine cells in the gut (magenta arrowheads) and intraspinal serotoninergic neurons (magenta empty arrowheads) (scale bar, 500 µm). (**D**) Illustrative images at 5 dpf of inner ear and lateral line neuromast hair cells, neuromast support cells and olfactory receptor neurons in either untreated (top), CuSO_4_-treated (middle) or BAPTA-treated (bottom) larvae. CuSO_4_ treatment selectively kills lateral line hair cells while BAPTA does not affect the overall morphology and organization of analyzed cells. HCs, hair cells (arrowheads, lateral line HCs; empty arrowheads, inner ear HCs from the 3 semi-circular canals); NECs, neuroepithelial cells (arrows); SCs, neuromast support cells (scale bars: 50 µm for inner ear region, 10 µm for all others). (**E**) Representative examples of trajectories of individual 5 dpf larvae with or without intact lateral line hair cells during 5 minutes. (**F**) Quantification of hair cells in the inner ear (N = 1 anterior macula or average over 3 cristae) or neuromasts (averaged over 4 neuromasts) in untreated (grey bars) versus CuSO_4_-(green bars) or BAPTA-treated (pale green bars) in 5-dpf larvae (t tests show significant loss of neuromast hair cells after treatment by CuSO_4_ with ****p = 1x10^-5^). (**G**) Treated larvae are able to locomote and perform escapes (N = 12 larvae per condition; t-tests show **p_CuSO4_ = 0.005 and **p_BAPTA_ = 0.001 compared to untreated siblings). (**H-J**) CuSO_4_-treated larvae (top panels, N = 54 untreated versus 55 treated siblings) exhibit drastic decrease in time spent swimming (**H**, Mann-Whitney test: ****p = 3x10^-13^), bout frequency (**I**, Mann-Whitney test: ****p = 3x10^-8^) and speed (**J**, Mann-Whitney test: ****p = 1x10^-10^). BAPTA-treated larvae (bottom panels, N = 39 untreated versus 38 treated siblings) show similar effects with decreased time spent swimming (**H**, Mann-Whitney test: ****p = 1x10^-9^), bout frequency (**I**, Mann-Whitney test: **p = 6x10^-3^) and speed (**J**, Mann-Whitney test: ****p = 5x10^-5^).

To test whether the modulation of spontaneous swimming relies on the mechanosensory-dependent activity of neuromast hair cells, we next impaired mechanoelectrical transduction only using BAPTA. BAPTA-treated larvae swam drastically less often, in a similar fashion than larvae lacking entirely their lateral line (Fig. 4E, and Movie S9), observed as an 81 % decrease in time spent swimming (Fig. 4H), a 32 % decrease in bout frequency (Fig. 4I) and a 50 % decrease in speed (Fig. 4J). Following the time course of tip link regeneration, mechanoelectrical transduction starts recovering after BAPTA washout within 30 minutes (*32*). Consistently, we observed a partial rescue of spontaneous locomotion in BAPTA-treated larvae 60 minutes after washout with aCSF (Fig. S2). Taken together, these results indicate that the spontaneous mechanosensory-dependent activity of lateral line hair cells can set the basal rate and speed of spontaneous swimming in zebrafish larvae.

### Zebrafish larvae rely on functional lateral line hair cells to respond to environmental serotonin

Finally, we tested the hypothesis that chemical activation by serotonin of lateral line hair cells contributes to the chemosensory behavior of larval zebrafish. We analyzed spontaneous swimming of larvae with intact or impaired lateral line navigating either in control or serotonin solution using the same conditions as above. We found that effects of serotonin on spontaneous locomotion were completely abolished in larvae treated with CuSO_4_ (Fig. 5A-B), suggesting that these effects are mediated by lateral line hair cells. As we could not rule out that this ototoxic agent may disrupt other cell types, we carried out the laser ablation of the posterior lateral line nerve to remove inputs from all neuromasts in the lateral line of the trunk. As a control to estimate side-effects of the manipulation itself on larval behavior, we performed a sham ablation away and parallel to the lateral line nerve (Fig. 5C, see Methods). Remarkably, the partial ablation of the lateral line was sufficient to abolish the effects of serotonin on time spent swimming (Fig. 5D) and to drastically reduce its effects on locomotor speed (Fig. 5E) compared to sham ablation.

**Fig. 5.**
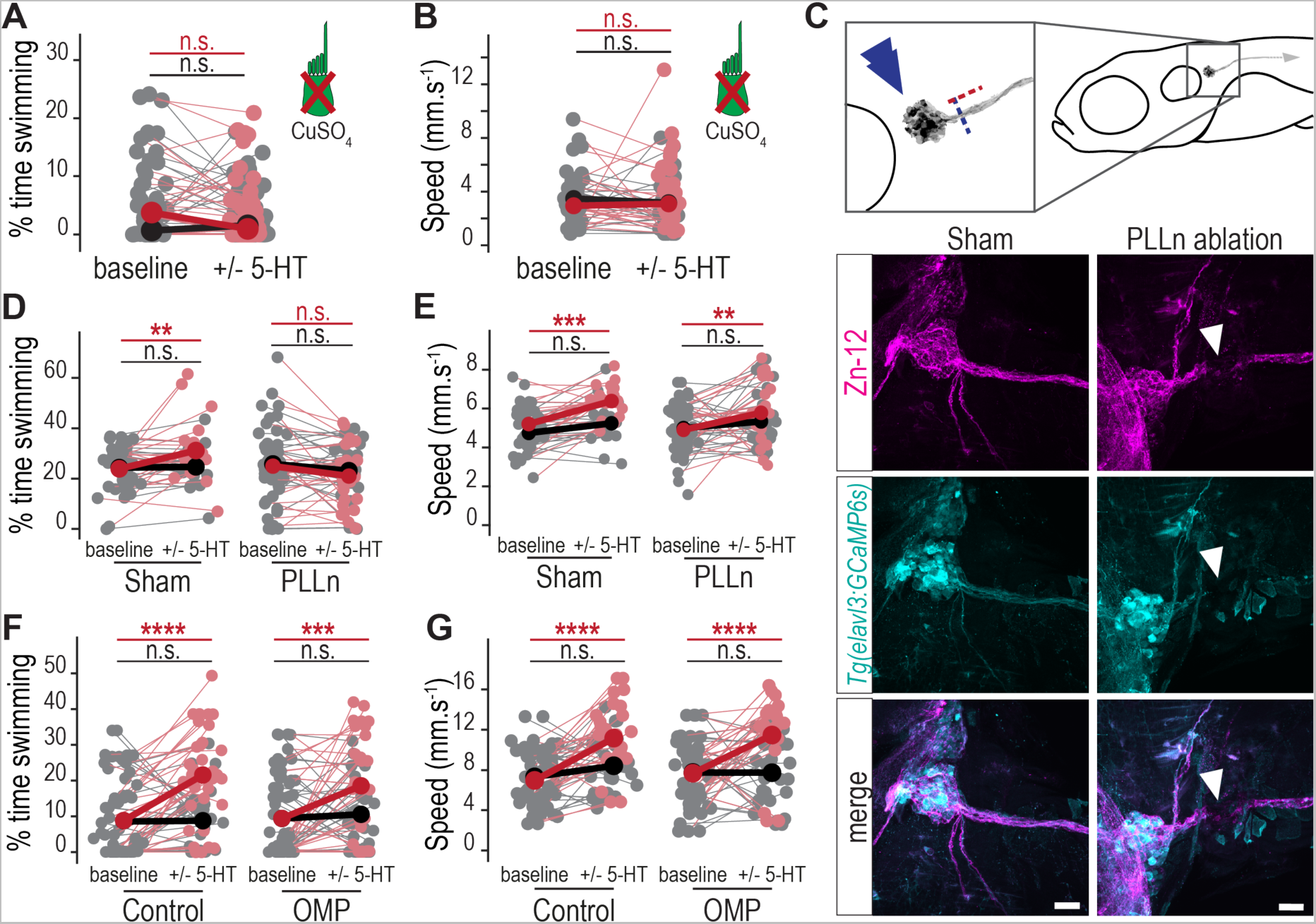
Zebrafish larvae rely on functional lateral line hair cells to respond to environmental serotonin. (**A-B**) CuSO_4_-treated larvae exposed to serotonin (magenta, N = 28 larvae) do not show increased swimming (**A**) and speed (**B**) from baseline, similar to siblings in control solution (grey, N = 31 larvae) (type II Wald chi-square tests indicate no significant interaction effect between time and condition for percentage of time swimming ^n.s.^p = 0.38, and speed ^n.s.^p = 0.72; statistics on the graph denote the results of post-hoc tests using Tukey’s method for the non-significant variation between baseline and test recordings). (**C**) The posterior lateral line nerve (PLLn) projects caudally from the posterior lateral line ganglion (blue double arrowheads) in 5 dpf *Tg(elavl3:GCaMP6s)* larvae. Laser ablation was performed either parallel and away from the nerve (red dotted line, sham controls) or across the nerve (blue dotted line, PLLn-ablated siblings). Illustrative images of co-immunohistochemistry for GCaMP (cyan) and the marker of neuronal cell surface Zn-12 (magenta) show the cut nerve (white arrowhead) in PLLn-ablated fish versus intact in sham siblings. Scale bars, 20 µm. (**D-E**) Laser ablation of the posterior lateral line nerve abolished the effect of serotonin on time spent swimming (**D**) and reduced its effect on speed (**E**) (N = 29 versus 24 larvae in control versus serotonin solution following ablation – “PLL” - compared to N = 19 versus 21 larvae in control versus serotonin solution in sham siblings – “Sham”; type II Wald chi-square tests indicate a significant interaction effect between time, condition and treatment for percentage of time swimming *p = 0.044; statistics on the left side of the graph indicate the significance of post-hoc tests using Tukey’s method for the variation between baseline and test condition in control – black – or serotonin – red – solution). (**F-G**) 5 dpf *Tg(ompb:GAL4; UAS:nitroreductase-tagRFP)* larvae following chemogenetic ablation of olfactory sensory neurons (N = 33 larvae in control solution – grey - versus 31 in serotonin solution - pink) show enhanced swimming upon exposure to serotonin similar to sham siblings (N = 33 larvae in control solution – grey - versus 33 in serotonin solution - pink) with increased time spent swimming (**F**) and speed (**G**) (type II Wald chi-square tests indicate significant interaction between time and condition but not treatment, confirming that olfactory ablation does not impact effects of serotonin on time spent swimming^5^; statistics on the graphs denote the results of post-hoc tests using Tukey’s method for the significant variation between baseline and test condition in control and serotonin solution).

As previously suggested, the olfactory receptor neurons could also be involved in carrying response to chemical cues in the surrounding water (*33*). To investigate the putative contribution of olfaction in the response to serotonin, we chemogenetically ablated olfactory sensory neurons in *Tg(ompb:GAL4;UAS:nitroreductase-mCherry)* larvae expressing the bacterial nitroreductase enzyme in *ompb*-positive olfactory sensory neurons using bath-application of the ligand metronidazole (*34*) prior to testing behavior (see Methods). Upon serotonin exposure, larval zebrafish deprived of olfactory receptor neurons (‘OMP’) exhibited, in a similar fashion than control siblings, a significant increase in time spent swimming (Fig. 5F) and speed (Fig. 5G, suggesting that the behavioral response to serotonin is not mediated by the olfactory system.

Together, these results indicate that zebrafish larvae rely on functional lateral line hair cells to elicit aversive behavior upon exposure to environmental serotonin.

## Discussion

We make here two major discoveries on the properties and functions of the lateral line in zebrafish. First, we discover that lateral line hair cells express a large repertoire of chemoreceptors known to respond to a diversity of chemicals such as serotonin, pheromones, amino acids, and physicochemical parameters like pH, temperature, and osmolarity. These results and our observation that some of these chemoreceptors are also detected in structures of the inner ear corroborate previous findings from recent hair cell transcriptomes revealing olfactory and purinergic receptors as well as TRP channels in larval (*35*) and adult (*36*) zebrafish, and in the mammalian cochlea (*8–13, 37*). Interestingly, many of these receptors have been identified in paddlefish lateral line (*38*) and in choanoflagellates (*39*), the closest unicellular living relatives of metazoan hair cells, pointing towards the conservation of these polymodal sensory functions among hair cells from a multisensory ancestor.

Second, we find that chemical compounds in the environment, such as serotonin, can activate lateral line hair cells and modulate their mechanosensory-driven activity, consequently impacting locomotion rate and speed. Our data indicates that, in the absence of external turbulent flow, the spontaneous mechanosensory activity of lateral line hair cells sets the basal level of spontaneous locomotion, a result that corroborates recent findings (*31*). In support, we observe a remarkable similarity between the distributions of previously-published inter spike intervals (ISIs) of afferent neurons (*40*) and interbout intervals (IBIs) we measured here. Our findings suggest that intrinsic mechanosensory-driven activity of lateral line hair cells could drive the initiation of bouts, likely via afferent neurons providing an excitatory drive to reticulospinal neurons in the hindbrain (*41*) that, in turn, send descending commands to spinal circuits to initiate movement. When confronted to water turbulences, our findings also suggest that lateral line hair cells may integrate both chemical and mechanical cues in order to complement the olfactory system for detecting the source of odors in a plume (*42, 43*), a fundamental process underlying complex behaviors such as feeding, mating, or communication between conspecifics (*44, 45*). In agreement with this hypothesis, a pioneering work on chemotaxis in sharks has shown that the lateral line is critical to track the trail of small-scale water turbulences as well as to pinpoint the sources of an odor in a plume (*46*). The discovery of chemoreception on the body via the lateral line also recalls the intriguing finding of chemoreception from the octopus’s arm (*47*). Further studies are necessary to investigate how complex chemical gradients are integrated with hydrostatic turbulences by lateral line hair cells in order to precisely locate odor sources in a rich environment.

In addition, the activation of lateral line hair cells by environmental serotonin is associated with qualitative changes in navigation, leading to avoidance and suggesting that chemosensory functions of lateral line participate in the implementation of elaborate behaviors. Recent studies have highlighted the diversity of zebrafish behavioral repertoire (*48, 49*) and demonstrated how complex behaviors, such as chemotaxis, involve recurrent sequences of simple maneuvers (*49*). With the development of novel algorithms for segmenting behavioral action sequences (*49*), future studies will be able to decipher temporal sequences elicited by chemical cues in the surrounding water.

Finally, we show that environmental serotonin is released by fish upon skin injury. The concentration of serotonin we detected in our conditions (nM range) in the skin extract of adult zebrafish falls in the range of EC50 known for receptors of the 5HTr3 family (*50*) (nM range in humans) and is thus sufficient to activate receptors expressed in neuromast hair cells. Our behavioral analysis reveals that serotonin elicits avoidance turns and could therefore act as an environmental cue of potential hazardous situations and predators by informing of the presence of injured conspecifics.

## Funding

European Research Council (ERC) Starting Grants “Optoloco” ERC-StG-311673 (CW) New York Stem Cell Foundation (NYSCF) Robertson Award 2016 Grant #NYSCF-R-NI39 (CW)

Human Frontier Science Program (HFSP) Research Grants #RGP0063/2014 (CW), #RGP0063/2018 (CW, LK) and #RGP0063/2017 (CW)

Fondation Schlumberger pour l’Education et la Recherche FSER/2017 (CW).

French Ministry of Higher Education and Research doctoral fellowship (LD)

Fondation pour la Recherche Médicale (FRM-FDT20170437143) (LD) and Team funding (FRM- EQU202003010612) (CW)

Gordon and Betty Moore Foundation Symbiosis Investigator Award GBMF9205 (https://doi.org/10.37807/GBMF9205) (Judith Eisen, funding LD)

Second Investissements d’Avenir program LIGHT4DEAF ANR-15-RHUS-0001 and LabExLIFESENSES ANR-10-LABX-65 (NM)

LHW-376 Stiftung (NM)

Fondation pour l’Audition FPA IDA03 (NM)

Fondation Bettencourt-Schueller (FBS-don-0031) Research Grant (CW) U19

NIH Consortium Grant #1U19NS104653 (CW)

Foundation Schlumberger for Research & Education (FSER) (CW)

## Author contributions

Conceptualization: CW, LD

Methodology: LD, JR, LK, NM, FXL

Investigation: LD, JR

Visualization: LD, JR, OM, LK, FXL

Funding acquisition: CW, LD, NM

Supervision: CW

Writing – original draft: CW, LD

Writing – review & editing: CW, LD, JR, NM, LK, FXL

## Acknowledgments

We wish to thank Prof. Katie Kindt for her generous gift of constructs and transgenic lines, as well as Yannick Marie and the ICM sequencing platform for their critical assistance in completing the transcriptomic analysis. We also thank Prof. Judith Eisen and Prof. Karen Guillemin for their generous support in completing some of the critical analysis as well as laser ablation experiments at University of Oregon with funding from the Gordon and Betty Moore Foundation.

## Supplementary materials

### Data and materials availability

Transcriptome data of lateral line hair cells is summarized in Data S1. Statistical analysis is summarized in Data S2. All other data, custom analysis codes and scripts are available on https://github.com/wyartlab/Desban_Roussel_Neuromasts_2022

## Materials and Methods

### Animal care and ethics statement

Animal handling and protocols were carried out with the validation of the Paris Brain Institute (Institut du Cerveau, ICM) in agreement with the French National Ethics Committee (“Comité National de Réflexion Ethique sur l’Expérimentation Animale”, APAFIS # 2018071217081175) and European Communities Council Directive (2010/63/EU). Adult zebrafish were reared at a maximal density of 8 animals per liter in a 14/10 hour light/dark cycle environment at 28.5°C. Larval zebrafish were typically raised in petri dishes filled with system water following the same conditions in terms of temperature and lighting as for adults. Experiments were performed at 20°C on animals aged between 3 and 5 days post fertilization (dpf) as described in each experimental protocol.

### Generation of transgenic lines

Transgenic lines used in this study are listed in the Table 1. To generate *Tg(myo6b:LifeAct-TagRFP,cryaa:mCherry)^icm33^* and *Tg(myo6b:eGFP)^icm29^*, Tol2 vectors were synthesized using 3-way Gateway recombination-based cloning with p5E-myo6b (kind gift of Prof. K. Kindt, NIH, USA), pME-eGFP or pME-LifeAct-TagRFP, and p3E-poly(A) into pDest or pDest-cryaa:mCherry destination vectors. The resulting product was injected into one-cell stage AB eggs at 30 ng/µL with 35 ng/µL Tol2 transposase RNA.

**Table 1.**
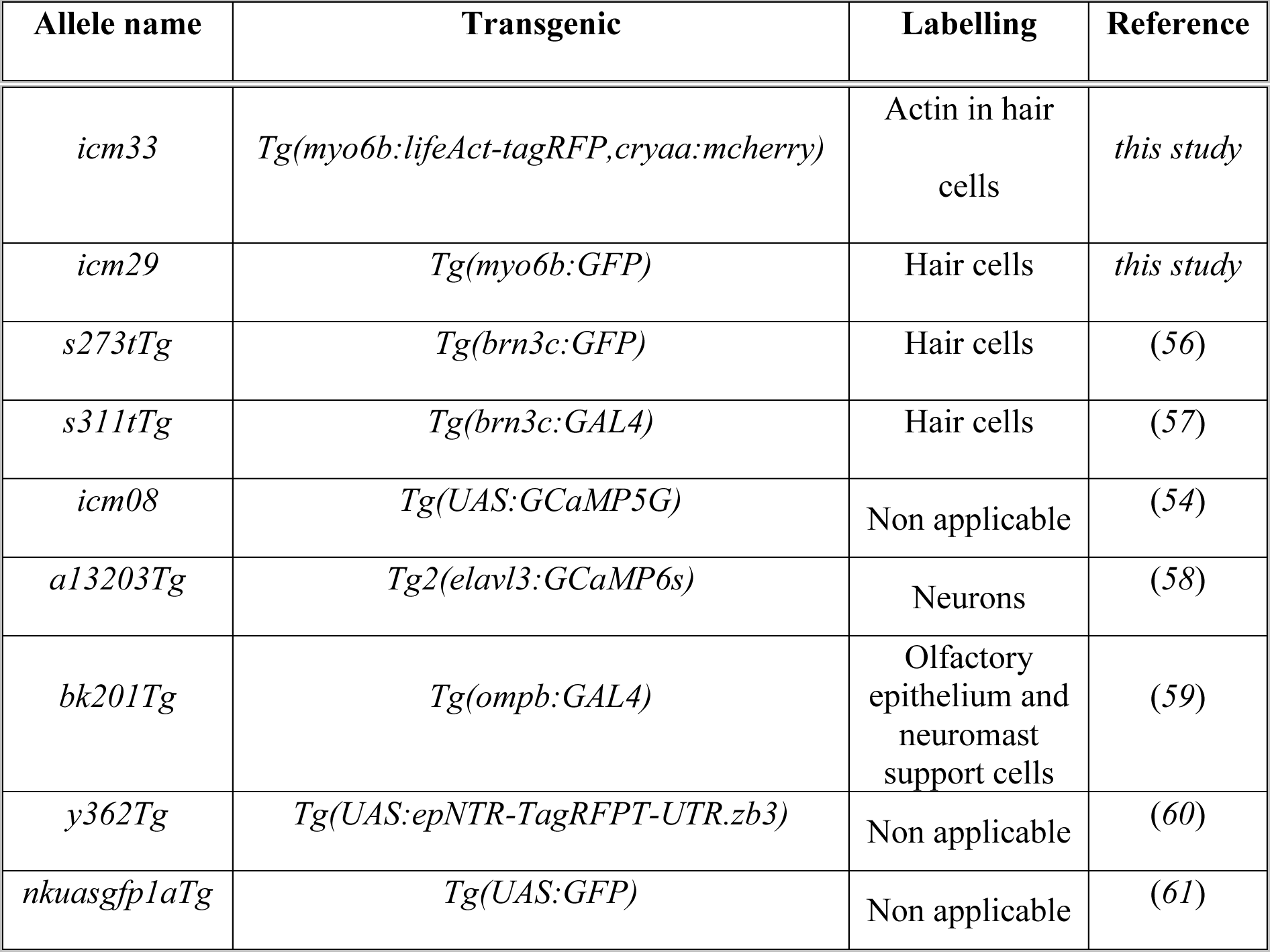
Alleles, specificity and origin of transgenic lines used in our study.

### Fluorescence activated cell sorting and validation

100 to 150 3-dpf zebrafish larvae were bathed in Yo-Pro-1 (4 µM, 30 min) (Invitrogen, Y3603), then washed 3 times in blue water to label lateral line hair cells. All larvae were then anaesthetized in 0.02 % (w/v) buffered 3-aminobenzoic acid methyl ester at pH = 7.0 (Tricaine; Sigma-Aldrich, A5040), immediately de-yolked in de-yolking buffer (55 mM NaCl, 1.8 mM KCl, 1.25 mM NaHCO_3_), and manually dissociated in FACSmax buffer (AMS Biotechnology, T200100) through a 40-µm sterile filter (Corning, #431750) following a previously established protocol (*62*). The resulting suspensions from Yo-Pro-1-labeled larvae were re-filtered through a 30-µm sterile filter prior to sorting on a BD INFLUX 500 Jazz sorter to isolate lateral line hair cells (between 3,000-15,000 cells/run, ∼ 0.2% of total input). The gating strategy was done by first selecting a homogenous population using both Side Scatter (SSC) and Forward Scatter (FSC). Singlets were then isolated based on FSC and trigger pulse width dispersion and living cells within the singlet population were selected based on UV absorption. Finally, the selection of the population of interested was based on green fluorescence after comparison with green-negative sample. In each sorting run, we isolated separate populations of putative lateral line hair cells and an assortment of Yo-Pro-1-negative cell types as a reference population, referred to as “Yo-Pro-1+” and “dark” cells respectively. Cells were sorted directly into lysis buffer and total RNA was quickly extracted using the RNAeasy Micro kit (Qiagen, #74004) for both qRT-PCR validation and RNAseq library preparation. Sorting runs were either entirely diverted to validation or library preparation as cell yield was insufficient to perform both analyses on the same sample. To verify that we isolated a highly-enriched population of lateral line hair cells prior to library preparation, several sorting runs were first diverted to qRT-PCR analysis. Total RNA was converted into cDNA using the SuperScript VILO cDNA synthesis kit (Thermo Fisher, #11754050) and relative levels of expression of a panel of diagnostic transcripts, including known genes specific of lateral line hair cells were compared between ‘Yo-Pro-1+’ and ‘dark’ pools using SYBR Green Chemistry (Thermo Fisher).

### Library preparation and total RNA sequencing

Once our approach validated by qRT-PCR, five additional sorting runs were performed. Total RNA from both ‘dark’ and ‘Yo-Pro-1+’ fractions extracted, and fragmented, tagged cDNA synthesized using the KAPA mRNA HyperPrep kit (Roche). The prepared libraries were sequenced on an Illumina NextSeq 500 using High Output Flowcell Cartridge from the NextSeq 500/550 Output v2 kit (75 cycles, up to 400 million reads, Illumina).

### Fluorescent immunohistochemistry of chemoreceptors

We applied to 5 dpf *Tg(brn3c:GFP)* transgenic larvae a similar protocol than previously described (*51*). Membrane GFP labelling was revealed using anti-GFP chicken primary antibody (1/500, Abcam, ab13970) and goat anti-chicken Fluor Alexa 488 secondary antibody (1/500, Invitrogen, A11039). The subcellular localization of the candidates 5HTr3a, Grp, Trpm4 and Trpa1 was revealed using rabbit IgG anti-5HTr3a (1/500, Abcam, ab13897), rabbit IgG anti-TRPM4 (1/200, Thermo Fisher, PA5-34283), rabbit IgG anti-GRP (1/200, Abcam, ab22623), and rabbit IgG anti-TRPA1 (1/200, Abcam, ab58844) primary antibodies respectively, and the goat anti-rabbit Fluor Alexa 568 secondary antibody (1/500, Invitrogen, #A11011). To obtain the labeling of Trpm4 in the inner ear, an additional step of antigen retrieval was performed prior to blocking using a 10-minute reaction with 10 µg/mL of Proteinase K in TE buffer (pH 8.0) at + 37 °C. Samples were mounted in 1.5% low-melting point agarose in glass-bottom dishes (MatTek, #P50G-1.5-14-F) and quickly imaged using Leica 40X and 63X oil-immersion objectives on a SP8 X white laser Leica inverted confocal microscope using Alexa Fluor 488 and 568 emission spectra to parameter hybrid detectors.

### ELISA

To detect and measure the concentration of serotonin in conditioned fish medium, we used the serotonin ELISA kit 96-strip-wells (My BioSource, MBS288208). The serotonin standard ranged from 50 ng/mL (284 nM) to 0.78 ng/mL (4.44 nM). Wells were filled with 50 µL of sample, standard or sample diluent (zero standard) before adding 50 µL of Detection Reagent A working solution. The plate was incubated 1 h at 37°C without agitation. Wells were then emptied by reversal and washed three times with 400 µL of wash buffer with 1 min of rest between each wash. 100 µL of Detection Reagent B working solution was added to each well prior to incubation for 45 min at 37°C without agitation. The incubation was followed by five washes. Finally, 90 µL of substrate solution was added to each well prior to incubation for 20 min at 37°C without agitation, away from light. The reaction was stopped by adding 50 µL of stop solution and results were read immediately on a 96-well reader SpectraMax M4 set to 450 nm. For analysis, the standard regression curve was first plotted as the log(target antigen concentration) as a function of the log(mean standard optical density) and fit to a linear regression. For samples with measured optical density above the control, the concentration in serotonin was estimated using a linear regression.

All reagents were allowed to reach room temperature prior to the experiment, all standards and controls were duplicated, and samples triplicated as three independent biological replicates.

### Whole-fish and skin extract preparation

Skin extract was prepared following a previously described protocol (*23*). Briefly, ∼ 1.5 year-old fish were quickly euthanized on ice and fifteen shallow cuts were done on each side of the trunk. For ‘Skin 1’ and ‘Skin 2’ skin extract, one or two lacerated fish, respectively, were transferred into a conical tube containing 2 mL of E3 medium and placed on ice. After 5 minutes, the E3 medium was retrieved from each tube and filtered using a 0.22-µm syringe filter (Sartorius, #16532) and a 1-mL syringe (Terumo, SS+01H1). Control ‘ctrl’ samples were obtained following the same procedure except that fish were left intact in the tube. The obtained skin extract was stored at 4°C prior ELISA testing during the same day, then kept at -20°C for later use. For whole-fish extracts, either single ∼1 month-old juveniles or ∼1.5 year-old adult zebrafish were quickly euthanized on ice prior to grinding using a 1-mL or 10-mL syringe in 0.5 or 5 mL of E3 medium, respectively. Obtained samples were then filtered using the same 0.22-µm filter and 1-mL syringe.

### Behavioral recordings

Zebrafish larvae were reared at density of 20 animals per 94 x 16 mm petri dish filled with 30 to 50 mL of control solution until 5 dpf. At 5 dpf, larvae were transferred to rectangular swim arenas (140 x 10 x 4 mm) filled with 4 mL of solution atop a white illumination panel where their spontaneous swimming was recorded for 5 or 10 minutes at 25 Hz using a ViewWorks camera (BASLER, acA2040-180km, https://www.baslerweb.com) controlled by the Hiris software (R&D Vision, https://www.rd-vision.com/r-d-vision-eng). Behavior of larvae following laser ablations of the posterior lateral line nerve were recorded in the same conditions but atop an infra-red illumination panel using an Edmund Optics camera (UI-3370SE USB 3.1) equipped with an infra-red filter and controlled by the uEyeCockpit software (iDS, https://www.ids-imaging.us/ids-software-suite.html). The first experiments were conducted in system water, and later in E3 medium. As we did not observe any significant change in locomotion of larvae in system water or E3 medium, nor across different days of experiment, we pooled the data acquired under these different conditions from different clutches. We referred to solutions devoid of drugs as “control solutions”. To test the role of serotonin (5-HT) in the modulation of spontaneous swimming, the basal level of locomotion of each larva was first evaluated in control solution. After 10 minutes, larvae were transferred into a second swim arena with either control or homogenous 5-HT (250 µM) fresh solutions and a second measure was taken to compare within and between individuals. Data in Fig. 2D-G was pooled from 5 different days of experiment (2 in system water, 3 in E3 medium). To assess the effect of local application of 5-HT, we pipetted 200 µL of 5 mM 5-HT solution in the central region restricted by custom combs in prefilled arenas. Individual larvae were then transferred at the extremity of each arena prior to removing the combs and starting the recording for 5 minutes. Data in Fig. 2I-K was pooled from 5 different days of experiment all performed in E3 medium. To test the role of lateral line hair cells in basal exploration, we compared spontaneous swimming of untreated versus CuSO_4_- or BAPTA-treated siblings according to conditions specified below (section ‘Chemical ablations using CuSO_4_ and BAPTA treatments and validation of treatment specificity’). For CuSO_4_, we pooled data from 4 days of experiments (2 performed in system water and 2 performed in E3 medium, Fig. 4H-J). For BAPTA, we pooled data from 3 days of experiment (1 performed in system water and 2 performed in E3 medium, Fig. 4H-J). To test the role of lateral line hair cells in the behavioral response to environmental serotonin, we compared spontaneous swimming of larvae in control solution versus homogenous 5-HT. We pooled data from 4 days of experiment for CuSO_4_-treated larvae (2 performed in system water and 2 in E3 medium, Fig. 5A-B), from 6 days of experiment for PLL-ablated larvae (all performed in E3 medium, Fig. 5D-E) and from 3 days of experiment for OMP-ablated larvae (all performed in E3 medium, Fig. 5F-G). The pH of all solutions was measured and buffered to be between 7,0 and 7,4.

### Simulation of the 5-HT gradient in the behavioral experiments

The concentration in the bath at any time is obtained from the numerical resolution of the unidimensional diffusion equation:

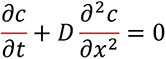

where *D* = 5.4·10^-10^ m^2^/s is the diffusivity of serotonin in water (*52*). The software Comsol (Finite Elements Methods) is used for the numerical simulations. A 1D mesh of 10000 elements discretizes a domain of length *L* = 14 cm. The initial condition is taken as *c*(0) = 1.46x10^-3^ mol/L for *x* ∈ [6.15; 7.85] cm and *c*(0) = 0 mol/L for *x* ∈ [0; 6.15] cm and *x* ∈ [7.75; 14] cm. To estimate these quantities, we assumed that during the injection process, 200 µL of solution at *c* = 5x10^-3^ mol/L were injected in a central region delimited by the comb of width of 1.7 cm, accordingly changing the local concentration to *c* = 1.46x10^-3^ mol/L due to dilution. We here neglected the change in the level of the bath induced by this injection of this small volume. A no-flux boundary condition is applied for *x* = 0 and *x* = 14 cm, which is expressed as ∂*c*/∂*x* = 0 for both coordinates. These results are represented in Fig. 2H for 5 min-long recordings, and a movie representing the evolution of the concentration in the bath as a function of time is provided as Movie S2. During the first 5 minutes of the process, the diffusion is too slow to significantly spread the serotonin far from the injection point.

### Tracking and analysis of behavior

Tracking of tail movements was performed to extract the timing of bouts, the position and head direction of the larvae and their tail angle using an improved version of the open source ZebraZoom algorithm (*27*) (www.zebrazoom.org). We estimated then the percentage of time spent swimming, bout frequency and bout speed for each larva using a custom MATLAB script. Forward swims and routine turns were distinguished using a cutoff of 30 degrees on the maximal bend amplitude of the tail angle.

### Calcium imaging

We monitored calcium activity of lateral line hair cells in live 3 dpf *Tg(brn3c:GAL4; UAS:GCaMP5G)* double transgenic larvae. Larvae were briefly anesthetized on ice in aCSF solution (119 mM NaCl, 26.2 mM NaHCO_3_, 2.5 mM KCl, 1 mM NaH_2_PO_4_, 1.3 mM MgCl_2_, 10 mM glucose, 2.5 mM CaCl_2_, oxygenated), transferred to a glass-bottom dish (MatTek, P50G-1.5-14-F) where they were immobilized under a nylon mesh (Smart Ephys, SHD-26GH/10) and paralyzed by injecting 0.5 nL of bungarotoxin at 0.5 mM (Tocris, #110.32-79-4) in the trunk musculature. Stimulations of individual neuromasts consisted in trials of 3 consecutive 1 s-long pressure-application of either control – aCSF alone with 10 µM Alexa 594 (Invitrogen, A10438) - or drug – aCSF with 10 µM Alexa 594 completed with 1 mM serotonin (Sigma-Aldrich, H9523) – solutions. Solutions were applied through a glass pipet (Sutter Instrument, BF-150-110-10) pulled with a micropipette horizontal puller (Sutter Instrument, P-1000) and controlled by a micromanipulator connected to a pneumatic picopump (4 psi, World Precision Instruments, SYS-PV820). Constant circulation of aCSF was maintained in the dish during the recording. Calcium imaging was performed using a Zeiss 63X water-immersion objective and an EMCCD Hamamatsu camera on a Zeiss Examiner microscope at 5 Hz for 210 s, allowing for 3 stimulations every 70 s and starting at 5 s in the recording. Images were acquired using Hiris software (R&D Vision) and reconstructed using Fiji (*53*). Manual selection of regions of interest (ROIs) – lateral line hair cell soma – and calcium imaging analysis were performed using a custom Matlab script (*51*). All image time series were registered in X and Y using a linear translation based on the first imaging frame to correct for motion artefacts and aligned to the stimulation time-course. We carefully excluded time points where drift in Z occurred during the 2 s-long time window following the pressure-application. Background noise, calculated in a region with no signal, was subtracted to every frame. Raw GCaMP signal was then extracted for every region of interest and the variation in signal ΔF / F in percent was calculated as F_GCaMP_ (t)*100/ F_0-GCaMP_ with F_GCaMP_ (t) the fluorescence at every time point and F_0-GCaMP_ the average basal level of fluorescence in a manually-selected quiet period. Each neuromast was tested twice: with aCSF control solution in Trial 1, followed by test solution in Trial 2. As explained in the results, Trial 1 investigated the mechano-sensory response of lateral line hair cells for a given orientation and enabled to distinguish responding from non-responding cells to a mechanical stimulus of given orientation by using a cutoff of 3-fold the ΔF / F standard deviation at baseline. In a pilot experiment, neuromasts were tested twice with aCSF solution, verifying that non-responding cells in Trial 1 remained silent when exposed to a stimulus with the same orientation in Trial 2 (Fig. 3D).

### Simulation of the 5-HT concentration in the calcium imaging experiments

In order to estimate the concentration of molecules at the surface of the neuromast during and after the puff application, we carried out 3D numerical simulations, applying finite-element method (Comsol Multiphysics, Transport of Diluted Species & Laminar Flow modules). The concentration of molecules *c* obeys the advection-diffusion law:

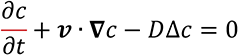

where *∇* is the differential operator and Δ is the Laplacian operator. Here *ν* is the fluid velocity vector obeying the incompressible Navier-Stokes equations:

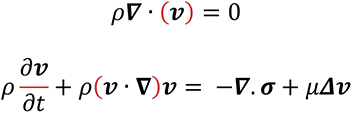

where *ρ* = 10^3^ kg/m is the water density and *σ* = –*pI* + *μ∇μ* + *μ*(∇*μ*)*^T^* is the total stress tensor, with *μ =* 10^-3^ Pa.s the water viscosity and *I* the identity matrix. Bold terms represent three dimensional vectors or tensors. The domain is a cylindrical container of height 0.25 mm and diameter 2 mm. The initial concentration *c*_0_ is set to 0. A flow of concentration *c*_puff_ passes through a spherical opening of diameter 3 μm, representing the injection pipette, with a flow rate of 3.6 nL/s during 1s, in the direction of the cell. The neuromast is represented by the union of a cone and an ellipsoid, of total height above the floor plate 45 μm. The neuromast surface is located at a distance of 30 μm from the pipette. A no slip boundary condition is imposed at the bottom wall and on the neuromast (*ν* = 0), and a complete impermeability is imposed against the passage of molecules (*n*·(*D*∇c + cν) = 0). The lateral walls as well as the upper wall of the cylindrical container are at imposed constant pressure *p*_0_, and we impose the absence of concentration gradients *n·∇*c = 0. Numerically, the transition between the injection process (which lasts 1s) and the post-injection phase (t > 1s) is achieved via a smoothing function:

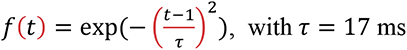

Taking advantage of the symmetry of the problem, we solved the equation in a half-domain discretized into 340’000 tetrahedral mesh elements. The simulations converged, and doubling the number of mesh elements led to similar results. The concentration distribution around the neuromast was found not to depend on the size of the container.

### Chemical ablations using CuSO_4_ and BAPTA treatments and validation of treatment specificity

To test the role of the lateral line system in spontaneous swimming, we used bath applications of waterborne copper sulfate (CuSO_4_, 50 µM in system water or 10 µM in E3 medium, for 2 hours at 28.5°C, Acros Organics, 10413465) to achieve cell-specific ablation or BAPTA (5 mM, 20 min in either system water or E3 medium at 28.5°C, Thermo Fisher, B1204) to disrupt mechanoelectrical transduction specifically. Treatments were stopped with several quick washes in the corresponding control solution (system water or E3). To verify the specificity of CuSO_4_ and BAPTA treatments, we assessed post-treatment hair-cell viability along with other chemosensory and superficial cell types exposed at the surface of 5 dpf larval zebrafish. We used the reporter lines *Tg(brn3c:GFP)* to label all hair cells from the inner ear and the lateral line, and *Tg(ompb:GAL4;UAS:GFP)* to label olfactory receptor neurons and neuromast support cells. Following a previously described protocol (*51*), we performed immunostainings for GFP using the anti-GFP chicken primary antibody (1/500, Abcam, ab13970) and goat anti-chicken Fluor Alexa 488 secondary antibody (1/500, Invitrogen, A110.39). We combined this immunohistochemistry with the one for serotonin (5-HT) using the rabbit IgG anti-5-HT (1/500, Sigma-Aldrich, 49545) and the goat anti-rabbit Fluor Alexa 568 (1/500, Invitrogen, A11011) to label neuroepithelial cells on the skin. Representative images displayed in Fig. 4 were taken from treatments performed in parallel on siblings in E3 medium.

### Manual estimation of the rate of escape responses in untreated versus treated 5 dpf larvae

*Tg(myo6b:GFP)* larvae were reared at density of 20 larvae in 30 mL of E3 medium in 94 x 16 mm petri dishes. All acousto-vestibular escapes were elicited on 5 dpf larvae using a behavioral setup previously described (*63*). Efficiency of the chemical treatment was assessed prior to any testing by verifying the absence of GFP-positive lateral line hair cells and that inner ear hair cells were intact. Larvae were then placed in round swim arenas (15 mm of diameter and 3 mm deep) filled with 1 mL of E3 medium atop an illumination panel using Light Emitting Diodes (LEDs). Escape responses were recorded using a ViewWorks camera (Basler acA2040-180km, https://www.baslerweb.com) controlled by the Hiris software (R&D Vision, https://www.rd-vision.com/r-d-vision-eng) for 1 second at 400 Hz. The escape stimulus consisted in an acoustic vibration occurring 200 ms after the onset of the recording. Each larva was subjected to 10 trials interspaced by 3 minutes of rest. We tested 12 CuSO_4_-treated larvae (10 µM in E3 medium, for 2 hours at 28.5°C) and 12 BAPTA-treated larvae (5 mM in E3 medium, 20 min at 28.5°C) along with 12 untreated siblings. We first quantified the ability of larval zebrafish to escape at least once across the 10 stimulations. Then we calculated the mean success rate of escape per condition as the number of elicited escapes over the 10 trials to assess the robustness of the response.

### Laser ablation of the posterior lateral line nerve

Transgenic *Tg(elavl3:GCaMP6s)* larvae were reared at density of 20 fish in 50 mL E3 medium in 94 x 16 mm petri dishes. At 5 dpf, larvae were anesthetized in 160 mg / L of 3-aminobenzoic acid methyl ester buffered at pH = 7.0 (Tricaine-S; Syndel) buffered at pH 7.0, then mounted laterally in 1.5 % agarose (Sigma-Aldrich, A6013) between slide and coverslip. Mounted larvae were placed under a 2-photon Ti:Sapphire microscope and fluorescent signal was detected through a 40X water-immersion objective with a laser tuned at 910 nm. After locating the posterior lateral line nerve emerging from the posterior lateral line ganglion, imaging mode was switched to line scan mode for 2 seconds during which laser power was focused on the two central lines, allowing effective laser ablation. Larvae were ablated in groups of 3 and on both sides by flipping the slide. After laser treatment, larvae were gently dismounted, transferred into fresh E3 medium and allowed at least 5 h of recovery prior to behavioral testing. All larvae were treated following the same protocol for the exception that control larvae were oriented laterally while ablated ones were oriented vertically resulting in sham versus posterior lateral line nerve ablations (Fig. 5C). We assessed efficiency of our laser ablations in larvae 5 h after treatment: following a previous protocol (*51*), we performed immunohistochemistry against GFP and Zn-12, a marker of neuronal cell surface, using the rabbit IgG anti-GFP (1/500, Invitrogen, A11122) and mouse IgG1 anti-Zn-12 (1/4000, Zebrafish International Resource Center Cat# zn-12, RRID:AB_10013761) primary antibodies, and goat anti-rabbit Alexa Fluor 488 (1/500, Invitrogen, A11008) and goat anti-mouse IgG1 Alexa Fluor 546 (1/500, Invitrogen, A21123) secondary antibodies.

### Chemogenetic ablation of olfactory receptor neurons

Double transgenic *Tg(ompb:GAL4; UAS:epNTR-tagRFPT-UTR.zb3)* and control single transgenic *Tg(ompb:GAL4)* siblings were manually dechorionated and transferred into 10 mM metronidazole (Sigma-Aldrich, M3761) dissolved in 0.1% DMSO (Sigma-Aldrich, 1010 D8418) for at least 54 hours starting at 2 dpf to induce the chemogenetic ablation of nitroreductase-expressing cells. Subsequently, 4 dpf larvae were then thoroughly washed 3 times with fresh E3 medium. After 24 hours of recovery, behavior recordings were performed on 5 dpf larvae comparing control siblings and ablated larvae – note that all animals were exposed the same way to metronidazole.

### Statistics

Non-parametric Mann-Whitney tests were used for hypothesis testing and ran on Matlab using the ‘ranksum’ function. To test the effect of serotonin on percentage of time spent swimming, bout frequency and bout speed, we used linear mixed-effect models coded in R (https://github.com/wyartlab/Desban_Roussel_Neuromasts_2022) after excluding outliers according to Cook’s distance greater than twice the amount of the cutoff value 4/n (for details about statistic methods and results, see Data S2). In all figures, * means that p < 0.05, ** means that p < 0.01, *** means that p < 0.001 and **** means that p < 0.0001 or equivalent after Bonferroni correction for multiple comparisons.

**Movie S1. Larvae swim faster and more often when exposed to serotonin.**

Typical examples illustrating the spontaneous swimming of two sibling larvae recorded in parallel at 25 Hz and played in real time in either control solution (top) or homogenous serotonin solution (250 µM, bottom). In presence of serotonin, larval zebrafish swam more often and faster as shown by increased bout frequency and bout speed. These examples depicted in Fig. 2C correspond to 30 s-long recording out of 5 min-long video matching the median behavior observed in the entire dataset. Numbers in the top right correspond to the bout numbers as tracked by ZebraZoom during the entire recording.

**Movie S2. The diffusion of a local source of serotonin leads to a sharp and local gradient that remains stable over 5-min-long recordings**

Evolution of the log of the concentration of serotonin (5-HT) in the bath following the pipetting of 200 µL of a solution at 5 mM in the central region delimited by custom-made combs. During a 5-minute-long recording, the diffusion is too slow to significantly spread serotonin far from the injection point, which results in the establishment of a steep gradient in the central area.

**Movie S3. The spontaneous swimming of 5 dpf larval zebrafish changes upon exposure to a local source of serotonin**

Typical examples of the spontaneous locomotion of two sibling larvae recorded in parallel at 25 Hz and played in real time in either control solution (top) or in a gradient of serotonin (bottom, 200 uL of 5 mM solution added to the central region of the well as delimited by the red rectangle). Encounter with serotonin induced more turns, and led zebrafish larvae to swim fast and away from the source. The example for the local gradient also depicted in Fig. 2C corresponds to 1-min-long recording out of a 5-min-long acquisition and matches the median behavior observed in the entire dataset. Numbers in the top right correspond to the bout numbers as tracked with the ZebraZoom algorithm during the recording.

**Movie S4. Stable fluorescence recording of neuromast hair cells upon hair bundle deflection by pressure-applied solutions**

A glass pipet filled with the test solution is positioned at the vicinity of the hair bundle of a neuromast. One stimulation consists in [5 seconds of rest – 1 second of puff – 64 seconds of rest] for a total of 70 seconds, i.e., 350 frames acquired at 5 Hz and played in real time. Each trial consists in 3 stimulations performed sequentially. During the 1s-long pressure-application, the hair bundle of hair cells in the vicinity is strongly deflected (left, brightfield) while the cell body remains in position, therefore enabling a stable recording. A perfusion system ensures the proper washout of solutions during the entire experiment as visualized using 10 µM of Alexa 594 dye in the test solution (right, red fluorescence displayed in grays), and allows for sequential tests.

**Movie S5. Serotonin rapidly diffuses following pressure-application at the vicinity of the neuromast hair bundle**

Evolution of the concentration of serotonin (5-HT) in the water surrounding the neuromast hair bundle protected by its cupula (black cone) during and 9 seconds after the 1-second pressure-application of a solution of concentration 1 mM in the glass pipet (black cylinder). Upon pressure-application, the maximal concentration reaching the cupula is estimated at 76 % of the original concentration, i.e., 760 µM, and rapidly decreases once the stimulus is off.

**Movie S6. A subset of hair cells shows reliable mechanical responses over consecutive trials with different pipets but same angle for pressure-application**

In a pilot study, neuromast hair cells were tested twice with aCSF during Trial 1 and Trial 2. The pressure-application of aCSF occurs for 1 second after 5 seconds of baseline during both trials (acquisition at 5 Hz, display in real time). Top, Regions of interest (ROIs 1 to 5) are drawn to outline hair cell soma and specifically follow the calcium activity in each cell within the neuromast. The same ROIs can be drawn from trial to trial to analyze activity of the same cells during consecutive trials. Responding hair cells are pointed out by red arrows. Bottom, calcium traces of the corresponding ROI.

**Movie S7. A subset of neuromast hair cells exhibit a strong and slow response to pressure-application of serotonin**

Neuromast hair cells were tested with control aCSF solution during Trial 1, then 1 mM serotonin (5-HT) during Trial 2. The pressure-application lasts 1 second and occurs after 5 seconds of rest during both trials. Fluorescence recordings were recorded and played at 5 Hz. Top, Regions of interest (ROIs, 1 to 3) are drawn to outline hair cell soma and specifically follow the calcium activity in each cell within the neuromast. ROI 2 responds largely to the mechanosensory stimulus but not serotonin. Conversely, ROI 3 does not respond to aCSF alone but to the application of 1 mM serotonin. The calcium transients in response to serotonin were slower both for on- and off-kinematics compared to aCSF. Bottom, calcium traces of the corresponding ROI.

**Movie S8. 5 dpf CuSO_4_-treated larvae exhibit severely impaired spontaneous locomotion compared to control siblings**

The spontaneous swimming of two sibling larvae, untreated (top) and CuSO_4_-treated (bottom), was recorded at 25 Hz during the same experiment (played in real time and shown in parallel). The treated larva exhibits severely reduced locomotion with rare and short swim bouts. These examples correspond to the ones depicted in Fig. 4E, of median behavior in the entire dataset, for 1 minute of recording in a 5-minute long video played in real time. Numbers in the top right correspond to the bout numbers as they were tracked with the ZebraZoom algorithm during the recording (https://zebrazoom.org/).

**Movie S9. 5 dpf BAPTA-treated larvae with impaired mechanoelectrical transduction exhibit similar reduced spontaneous locomotion than larvae with no lateral line hair cells**

Typical examples illustrating the spontaneous swimming of two sibling larvae (untreated (top) and BAPTA-treated (bottom)) recorded at 25 Hz during the same experiment, played in real time and shown in parallel. Compared to control siblings, the BAPTA-treated larva exhibits similar exploration deficits (with few locomotor bouts occurring at low speed) than larvae lacking lateral line hair cells after copper-sulfate treatment. This example corresponds to the one depicted in Fig. 4E with 1 min-long recording from the 5 min-long video and reflects the median behavior observed in the entire dataset. Numbers in the top right correspond to the bout numbers as they were tracked with the ZebraZoom algorithm during the recording (https://zebrazoom.org/).

**Data S1. Transcriptome dataset of Yo-Pro-1-positive versus –negative fractions averaged across five biological replicates**

**Data S2. Statistical analysis of behavioral data**

**Fig. S1.**
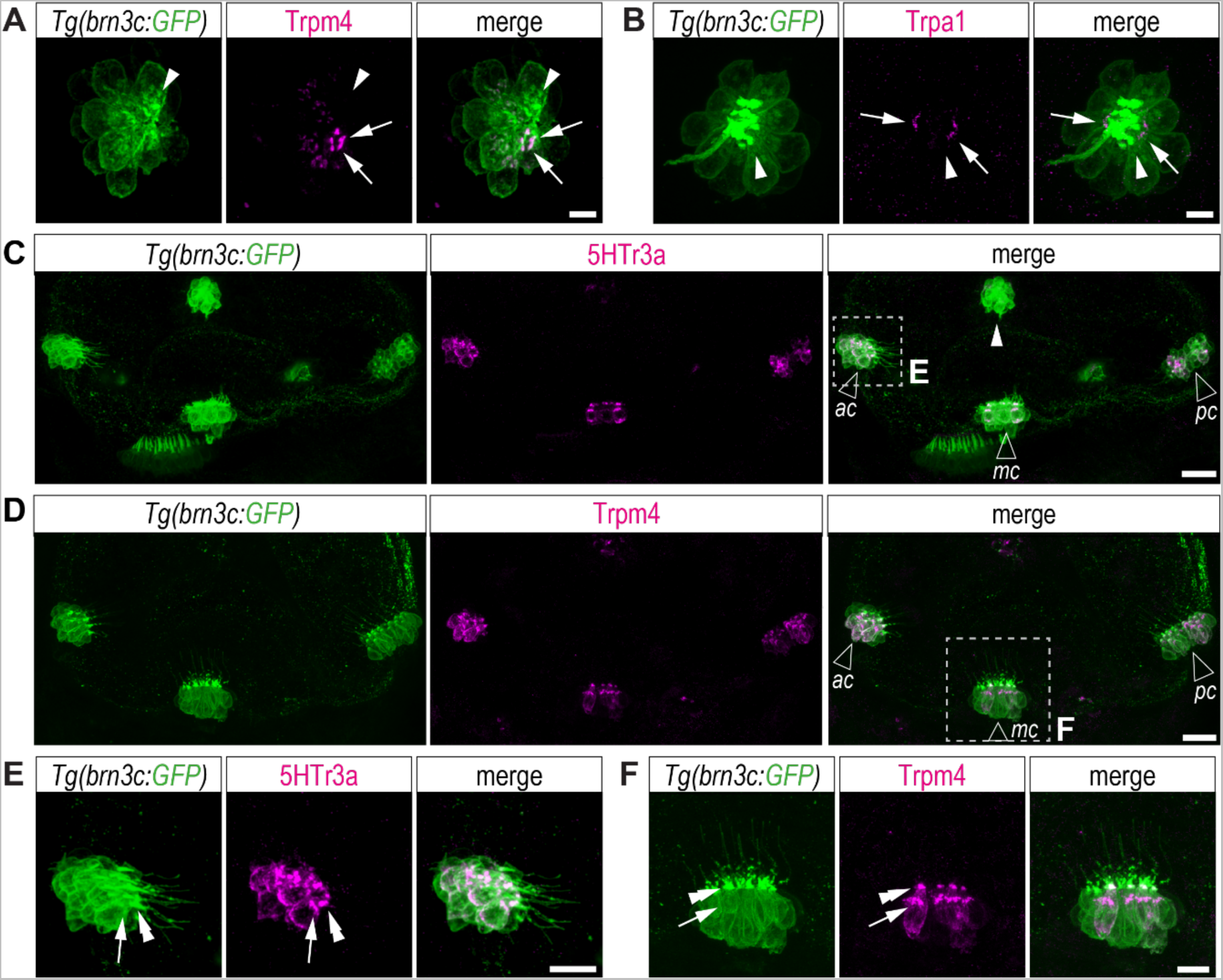
Chemoreceptors are found in a subset of hair cells in the lateral line and inner ear. Immunostainings reveal Trpm4 (**A**) and Trpa1 (**B**) in the subapical region (arrows) of a subset of GFP-positive lateral line hair cells at 5 dpf. Arrowheads, non-expressing cells. Scale bars, 5 µm. 5HTr3a (**C**) and Trpm4 (**D**) are also expressed in GFP-positive inner ear hair cells from the three cristae of the semi-circular canals (empty arrowheads; ac, anterior crista; mc, medial crista; pc, posterior crista). Scale bars, 20 µm. Close-up of regions depicted in **C** and **D** show the concentration of 5HTr3a (**E**) and Trpm4 (**F**) in the subapical (Golgi apparatus, arrows) and apical (cuticular plate, double arrowheads) regions in a fraction of inner ear hair cells from one crista. Scale bars, 10 µm.

**Fig. S2.**
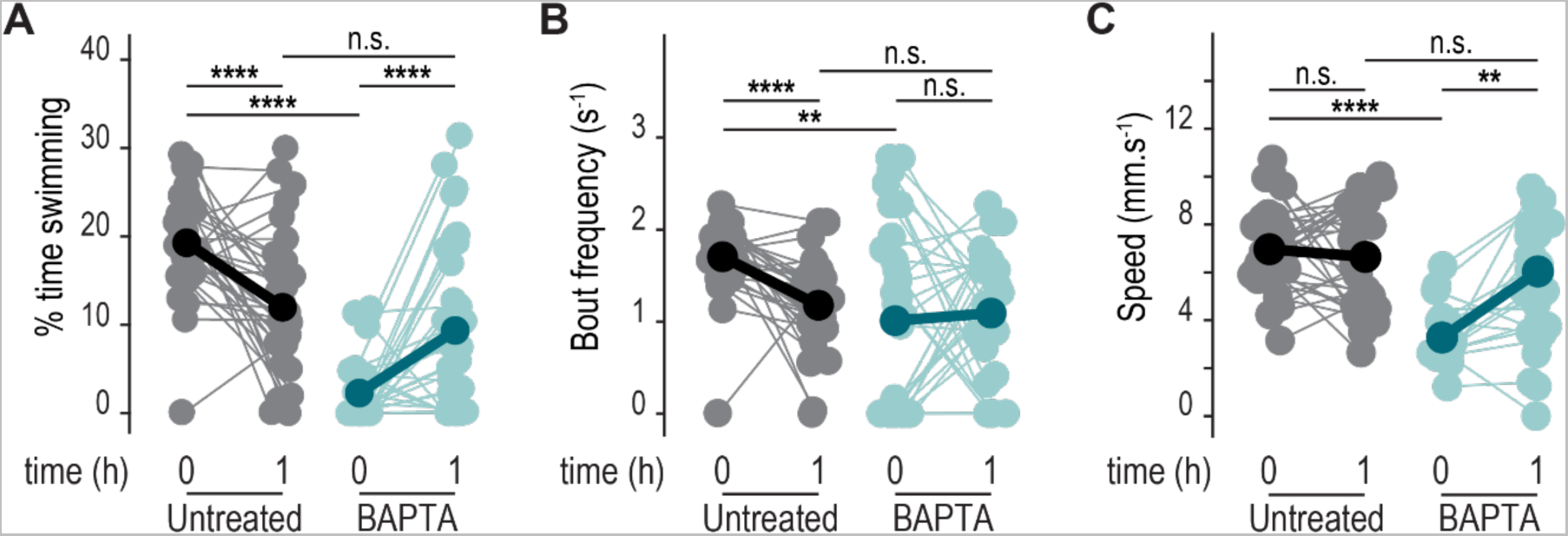
BAPTA-treated larvae show partial recovery within 1-hour post-washout. Right after washout (time “0”), BAPTA-treated larvae display strong reduction in exploratory behavior with significant reduction in time spent swimming (**A**, Mann-Whitney test: ****p = 1.9x10^-11^), bout frequency (**B**, Mann-Whitney test: **p = 2.9x10^-3^) and speed (**C**, Mann-Whitney test: ****p = 1.7x10^-8^) compared to untreated siblings (N = 33 treated versus 33 untreated animals). One hour after washout (time “1”), untreated larvae show habituation with less time spent swimming (**A**, repeated measure ANOVA: ****p = 7.2x10^-5^), reduced bout frequency (**B**, repeated measure ANOVA: ****p = 9.7x10^-6^) but constant speed (**C**, repeated measure ANOVA: ^n.s.^p = 0.44) while BAPTA-treated siblings show partial recovery with increase in time spent swimming (**A**, repeated measure ANOVA: ****p = 6.3x10^-5^), bout frequency (**B**, repeated measure ANOVA: ^n.s.^p = 0.72) and speed (**C**, repeated measure ANOVA: **p = 5.6x10^-3).^ One hour after BAPTA treatment, treated-larvae exhibit consequently similar kinematics than untreated siblings (Mann-Whitney tests: ^n.s.^p_swim_ = 0.10, ^n.s.^p_FQ_ = 0.97 and *p_speed_ = 0.07).

